# Serotonergic neural network related to behavioral inhibition system

**DOI:** 10.1101/2021.07.09.451787

**Authors:** Kazuho Kojima, Shigeki Hirano, Yasuyuki Kimura, Chie Seki, Yoko Ikoma, Keisuke Takahata, Takehito Ito, Keita Yokokawa, Hiroki Hashimoto, Kazunori Kawamura, Ming-Rong Zhang, Hiroshi Ito, Makoto Higuchi, Satoshi Kuwabara, Tetsuya Suhara, Makiko Yamada

## Abstract

*Rationale* The tendency to avoid punishment, called behavioral inhibition system, is an essential aspect of motivational behavior. Behavioral inhibition system is related to negative affect, such as anxiety, depression and pain, but its neural basis has not yet been clarified. *Objectives* To clarify the association between individual variations in behavioral inhibition system and brain 5-HT_2A_ receptor availability and specify which brain networks were involved in healthy male subjects, using [^18^F]altanserin positron emission tomography and resting-state functional magnetic resonance imaging. *Result*s Behavioral inhibition system score negatively correlated with 5-HT_2A_ receptor availability in anterior cingulate cortex. A statistical model indicated that the behavioral inhibition system score was associated with 5-HT_2A_ receptor availability, which was mediated by the functional connectivity between anterior cingulate cortex and left middle frontal gyrus, both of which involved in the cognitive control of negative information processing. *Conclusions* Individuals with high behavioral inhibition system displays low 5-HT_2A_ receptor availability in anterior cingulate cortex and this cognitive control network links with prefrontal-cingulate integrity. These findings have implications for underlying the serotonergic basis of physiologies in aversion.

## Introduction

The fundamental features of complex behavior have long been discussed as being categorizable into the approach to rewards and the avoidance of punishments. These two systems can be applied to account for personality and motivation (Davidson, 1994; Gray, 1982; Higgins, Roney, Crowe, & Hymes, 1994), positing that there are independent sensitivity in the respective systems. Gray provided a powerful theoretical framework that was rooted in behavioral psychology and neuroscience, called Reinforcement Sensitivity Theory (RST) (Gray, 1982). Gray proposes two systems together with an additional one: Behavioral Approach System (BAS), Behavioral Inhibition System (BIS), and Fight-Flight System (FFS) (see revised version of RST(Gray & McNaughton, 2000)). BAS corresponds to impulsivity, drug addiction, and attention deficit hyperactivity disorder, BIS is related to anxiety, depression, and pain, and FFS is fear at the psychological and psychiatric level (Bijttebier, Beck, Claes, & Vandereycken, 2009; P.J. Corr, 2002; P. J. Corr, 2004; Jensen, Ehde, & Day, 2016).

In contrast to BAS, however, only a handful of studies have investigated the neural basis of BIS. The trait sensitivity to aversive events was associated with increased gray matter volume in amygdala and hippocampus (Barros-Loscertales et al., 2006; Cherbuin et al., 2008) and decreased volume in orbitofrontal cortex (OFC) and precuneus (Fuentes et al., 2012). BIS variability was also associated with individual differences in the neural activities of dorsal anterior cingulate cortex (ACC), OFC, striatum, amygdala, and hippocampus during anticipation of aversive events, such as monetary loss, measured by functional magnetic resonance imaging (fMRI) (Beaver, Lawrence, Passamonti, & Calder, 2008; Kim, Yoon, Kim, & Hamann, 2015; Simon et al., 2010). A resting-state fMRI (rs-fMRI) study similarly found that BIS correlated negatively with regional homogeneity in amygdala and hippocampus (Hahn, Dresler, Pyka, Notebaert, & Fallgatter, 2013).

Meanwhile, a number of neuroimaging studies have investigated the neural responses to aversive stimuli such as signal pain, punishment and monetary loss. The core regions of these aversive anticipations are found in ACC, anterior insula, OFC, and amygdala (De Martino, Camerer, & Adolphs, 2010; Eisenberger, 2012; Hayes & Northoff, 2011; Kringelbach & Rolls, 2004; Nitschke, Sarinopoulos, Mackiewicz, Schaefer, & Davidson, 2006; Wrase et al., 2007). Congruent brain regions (ACC, OFC, amygdala) between aversive anticipation and individual variations in the sensitivity to aversive events leads to the notion that these regions are the hub for understanding the neural mechanisms of BIS.

The serotonergic system is supposed to be associated with behavioral inhibition in the human brain (Cloninger, 1987). Notably, serotonin 2A (5-HT_2A_) receptor has been reported to be intimately involved in the modulation of negative emotions, such as anxiety, depression, and pain (Baldwin & Rudge, 1995; Sommer, 2009). According to Gray’s concept of BIS, harm avoidance is characterized as excessive anxiety and fear, but the evidences for the role of serotonergic system in harm avoidance, so far remains inconclusive. For instance, harm avoidance and 5-HT_2A_ receptor availability showed a negative correlation in prefrontal cortex and left parietal cortex (Moresco et al., 2002), a positive correlation in dorsal prefrontal cortex (Baeken, Bossuyt, & De Raedt, 2014), or no significant regional correlation (Soloff, Price, Mason, Becker, & Meltzer, 2010). Although human positron emission tomography (PET) studies have shown that the 5-HT_2A_ receptors are highly and widely distributed in cortical regions (Savli et al., 2012), it remains unclear whether the individual variations in 5-HT_2A_ receptor availability are involved in the trait sensitivity to aversive events, i.e., BIS.

The aim of this study was to elucidate the neural and molecular mechanisms associated with individual variations in BIS. In this regard, the relationships among BIS, the 5-HT_2A_ receptor availability using PET and the brain functional connectivity measured by rs-fMRI were investigated. We first conducted a PET imaging study to explore which brain regions of 5-HT_2A_ receptor availability correlated with BIS in healthy volunteers. Then, we analyzed rs-fMRI data to detect functional connectivity showing correlation with local 5-HT_2A_ receptor availability and BIS. Finally, mediation analysis was conducted to elucidate the relationships among BIS, functional connectivity and the 5-HT_2A_ receptor availability.

## Materials and methods

### Participants

Sixteen healthy right-handed male subjects (age: 23.3 ± 2.9 years, mean ± standard deviation) were recruited. Two subjects were excluded due to incomplete data collection, and the data of fourteen participants (23.4 ± 2.9 years) were analyzed. All participants were free of current and past psychiatric or somatic disorders, and had no history of drug abuse. Each participant completed psychological testing and underwent both rs-fMRI and PET scans. All participants provided written informed consent before participating in the study, which was approved by the Ethics and Radiation Safety Committee of the National Institute of Radiological Sciences in accordance with the ethical standards laid down in the 1964 Declaration of Helsinki and its later amendments.

### Psychological measurement

To test Gray’s original theory, Sensitivity to Punishment and Sensitivity to Reward Questionnaire (SPSRQ) was developed by Torrubia (Torrubia, Ávila b, Moltó, & Caseras, 2001). This scale indicates good reliability and validity, and accurately expresses the essence of Gray’s theory (e.g., extraversion and neuroticism in expected directions). All participants completed the Japanese version of SPSRQ (Takahashi & Shigemasu, 2008). SPSRQ is a 48-item self-report measure that consists of two subscales, representing sensitivity to reward (SR) to measure impulsivity, i.e., BAS, and sensitivity to punishment (SP) to measure anxiety, i.e., BIS. Each item is scored on a 4-point Likert scale (1 = disagree, 4 = agree). SR and SP scores with higher scores indicating greater impulsivity and sensitivity to punishment, respectively. Participants also completed the Beck Hopelessness Scale (BHS(Beck, Weissman, Lester, & Trexler, 1974)) and State-Trait Anxiety Inventory (STAI (Spielberger, Gorsuch, Lushene, Vagg, & Jacob, 1983)) to measure the levels of depressive hopelessness and anxiety, respectively.

### PET acquisition and analysis

All subjects underwent a PET scan to measure regional 5-HT_2A_ receptor availability. A 90-min dynamic PET acquisition was performed after an injection of [^18^F]altanserin (190 ± 5.4 MBq with molar activity of 167 ± 77 GBq/μmol). The scan protocol consisted of 33 frames (10 s × 6, 20 s × 3, 1 min × 6, 3 min × 4, and 5 min × 14 frames). All of the PET scans were performed on an Eminence SET-3000 GCT/X PET scanner (Shimadzu; Kyoto, Japan) with a head fixation device to minimize head movement. Each PET scan was preceded by a transmission scan for attenuation correction using a ^137^Cs source. All PET images were reconstructed with the filtered back-projection method (Gaussian filter, kernel 5 mm; reconstructed in-plane resolution was 7.5 mm in full width at half maximum; voxel size: 2 × 2 × 2.6 mm) corrected for attenuation, randoms and scatter.

During the scans, arterial blood samples were obtained manually 33 times after radioligand injection to obtain arterial input function (Ishii et al., 2017). Each blood sample was centrifuged to obtain plasma and blood cell fractions, and the concentrations of radioactivity in whole blood and plasma were measured (Ishii et al., 2017). The fractions of the parent compound and its radiometabolites in plasma were determined using high-performance liquid chromatography from 6 samples of each subject (Ishii et al., 2017).

All PET images were spatially normalized to the standard anatomic orientation. First, head motion during the scans was corrected on the emission images after correction of attenuation using μ-maps that were realigned to each frame of the emission images (Wardak et al., 2010). Second, T1-weighted MR images were coregistered to the corresponding mean PET images. Third, the MR images were spatially normalized and segmented into gray matter, white matter, and cerebrospinal fluid using SPM8 (Wellcome Institute of Neurology, University College of London, UK). Finally, all PET images were spatially normalized to the standard anatomic orientation (Montreal Neurological Institute (MNI) 152 standard space; Montreal Neurological Institute; Montreal, QC, Canada) based on the transformation of the MR images.

Because the Logan analysis provided a good compromise between validity, sensitivity, and reliability of implementation (Price et al., 2001), the PET data were analyzed by Logan graphical method (Logan et al., 1990), which was applied across the 12-to 90-min integration intervals, and regional total distribution volume (*V*_T_) values were obtained. We used the cerebellum as reference brain region and estimated the nondisplaceable distribution volume (*V*_ND_). 5-HT_2A_ receptor availability was determined as binding potential (*BP*_P_) that was derived from the equation: *BP*_P_ = *V*_T_–*V*_ND_ (Innis et al., 2007). All kinetic analyses were performed using PMOD (version 3.6, PMOD Technologies Ltd., Zurich, Switzerland).

### Region-of-interest analysis

ACC, OFC and amygdala, which are involved in the sensitivity to aversive events (Barros-Loscertales et al., 2006; Beaver et al., 2008; Cherbuin et al., 2008; Eisenberger, 2012; Fuentes et al., 2012), were applied to ROI analyses. These brain regions were extracted by the Harvard-Oxford atlas using the CONN toolbox (version 17e, http://www.nitrc.org/projects/conn), averaged over right and left. Subsequently, ACC was divided into four segregated subregions, namely, subgenual ACC (sgACC), pregenual ACC (pgACC), anterior midcingulate cortex (aMCC) and posterior midcingulate cortex (pMCC) (Yeung, Botvinick, & Cohen, 2004). Subgenual ACC was substituted by subcallosal cortex in the atlas due to small volume (0.056 mm^3^). The volumes of each ACC subregion were 9.2 mm^3^ in subcallosal cortex, 6.5 mm^3^ in pgACC, 9.2 mm^3^ in aMCC, and 6.7 mm^3^ in pMCC.

### Resting-state fMRI acquisition and analysis

Each subject underwent a 6.8-minute rs-fMRI scan, performed with a Magnetom Verio 3.0 T MRI scanner (Siemens, Erlangen, Germany) equipped with a 32-channel head coil. During scanning, subjects were instructed to relax with their eyes open while gazing at a fixation cross. A single session acquired 3.8-mm thick, no gap, interleaved axial 33 slices (in-plane resolution: 3.75 × 3.75 mm) with a 30-degree angle relative to the AC-PC axis, using a T2*-sensitive single-shot EPI sequence with the following parameters: TR = 2000 ms, TE = 25 ms, flip angle = 90 degrees, matrix = 64 × 64. A high-resolution T1-weighted anatomical image using a magnetization prepared rapid acquisition gradient echo (MPRAGE) sequence (176 sagittal slices, resolution = 0.49 × 0.49 × 1.00 mm, no gap, TR = 2300 ms, TE = 1.95 ms, flip angle = 9 degrees, matrix = 512 × 512) was acquired for anatomical reference.

Data processing was performed using the CONN toolbox and SPM12 (Wellcome Institute of Neurology, University College of London, UK) working on Matlab version 8.4 (MathWorks, MA, USA). The first four volumes were discarded from analysis to account for magnetization saturation effects. Preprocessing comprised: 1) realignment and unwarping, 2) slice timing correction, 3) segmentation and normalization, 4) smoothing with a Gaussian kernel of 4 mm. To eliminate correlations caused by head motion and artifacts, we identified outlier time points in the motion parameters and global signal intensity using Artifact Detection Tools (ART), which includes the CONN toolbox. For each subject, we treated images as outliers if composite movement from a preceding image exceeded 0.2 mm, or if the global mean intensity was over 3 SDs from the mean image intensity for the entire resting scan. Based on the previous report ^62^, after removing outlier images, one subject whose total scan time of less than 3 minutes was excluded from the subsequent analyses.

After preprocessing, we conducted de-noising as follows: 1) linear regression of noise sources from white matter and cerebrospinal fluid by CompCor (component-based noise correction method) and from outliers by ART and from Friston 24 head motion parameter, 2) band-pass filtering of 0.009 – 0.1 Hz was used to pass the low frequency fluctuations of interest, 3) quadratic trends were removed. Global signal regression was not used to avoid potential false anticorrelations.

### Image analyses

#### 1. Association between SP and 5-HT_2A_ receptor availability

To test for the contribution of 5-HT_2A_ receptor availability to SP, we first conducted Spearman’s rank test between SP and the *BP*_P_ value of each ROI using GraphPad Prism (version 7, GraphPad Software, CA, USA). P-value less than 0.05 with false discovery rate (FDR) correction for multiple comparisons was considered significant.

#### 2. Relationship between 5-HT_2A_ receptor availability and functional connectivity

Regions of 5HT_2A_ receptor availability that correlated with SP were subsequently investigated by seed-based functional connectivity analysis modeling the *BP*_P_ values in the analogous regions, utilizing the CONN toolbox. This procedure may explore the functional connectivities that correlate with 5-HT_2A_ receptor availability of SP-related ROIs. The threshold was defined as a cluster-level threshold of p < 0.05, FDR-corrected with voxel-level threshold of p < 0.001, uncorrected for multiple comparisons. All reported coordinates were of MNI standard space.

#### 3. Functional connectivity related to SP

The correlation coefficient for each specified functional connectivity was extracted. Spearman’s rank test was performed between each extracted correlation coefficient and SP. P-value less than 0.05 was considered significant.

#### 4. Mediation analysis

Finally, for each functional connectivity that was significantly related to both 5-HT_2A_ receptor availability and SP, we performed mediation analysis to test whether the functional connectivity might be involved in the link between SP and 5-HT_2A_ receptor availability. The correlation coefficient of functional connectivity was included as a mediator. INDECT macro (Preacher & Hayes, 2008) with SPSS (version 24, IBM, NY, USA) were used. Bias-corrected and accelerated 95% confidence intervals based on 10000 bootstrap sampling were used to assess significance.

## Results

### Behavioral findings

The median SP and SR scores were 61 (interquartile range, 55 to 67.5) and 51 (interquartile range, 44 to 59), respectively. The median score of BHS was 6 (interquartile range, 4.5 to 10.5) and STAI was 41 (interquartile range, 34 to 54.5). SP scores correlated positively with BHS (r_s_ = 0.86, p < 0.0005, Spearman’s rank test) and at a marginally significant level with STAI (r_s_ = 0.52, p = 0.07).

### Regional 5-HT_2A_ receptor availability measured by [^18^F]altanserin PET

Figure 1 shows the 5-HT_2A_ receptor availability values (*BP*_P_) in ACC, OFC and amygdala which were selected from the premise that these regions were associated with aversive anticipation. Higher *BP*_P_ was measured in the ACC and OFC, while low *BP*_P_ was observed in the amygdala. The *BP*_P_ values in ACC (1.535 [1.376 to 1.773]), OFC (1.503 [1.41 to 1.633]) and amygdala (0.676 [0.581 to 0.762]) were comparable to that of healthy subjects in previous report (Savli et al., 2012).

**Figure 1.**
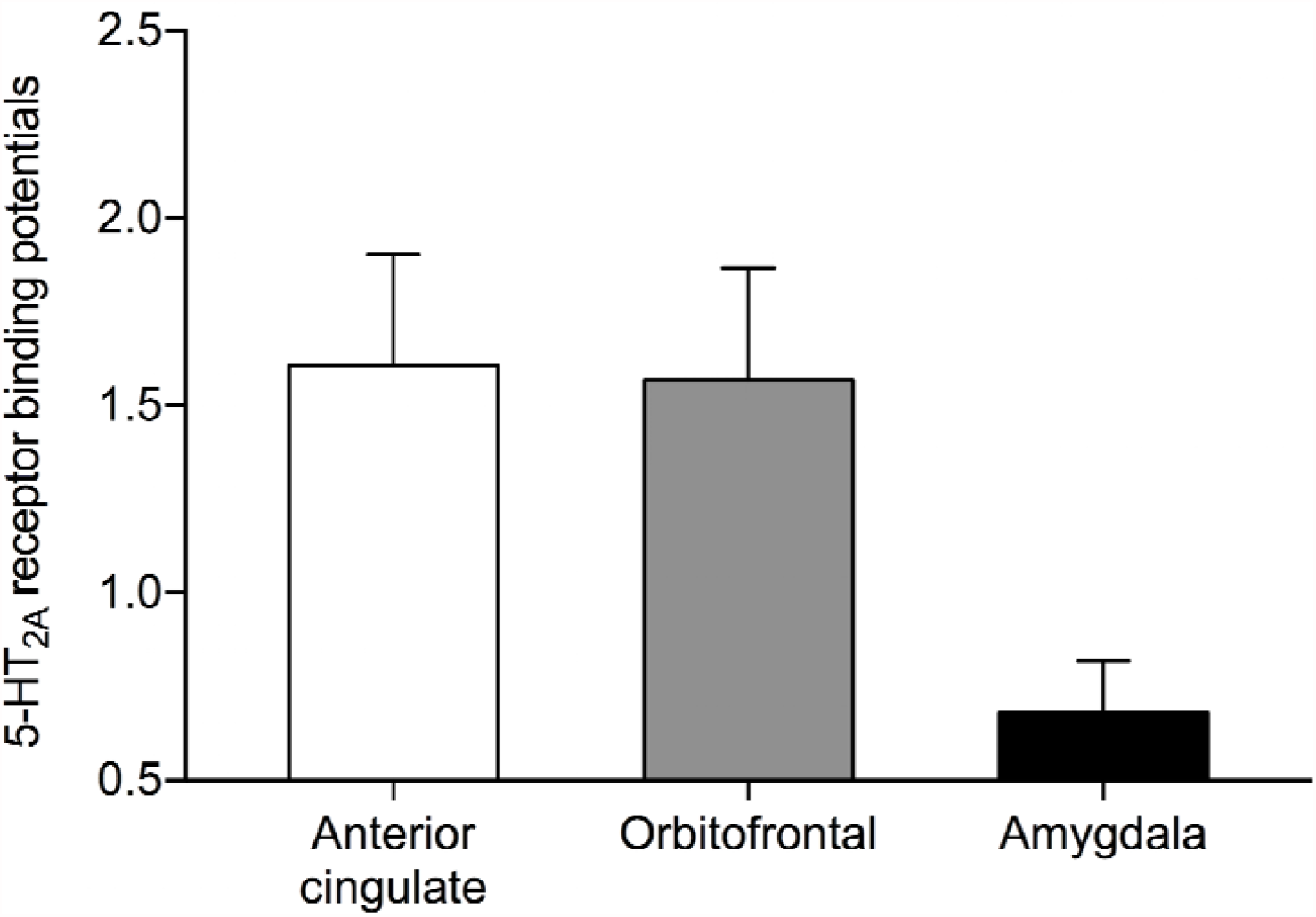
[^18^F]altanserin binding potentials of the limbic system. Bar graphs represent mean ± standard deviation.

### Association between behavioral inhibition system and 5-HT_2A_ receptor availability

The SP score was negatively correlated with the *BP*_P_ value of ACC (r_s_ = -0.66, p = 0.016, Spearman’s rank test with FDR correction). No correlations were found in OFC or amygdala (r_s_ = -0.47 and r_s_ = -0.57, respectively, both p > 0.05 with FDR correction).

We further examined the correlations between SP and the *BP*_P_ values in the functional subdivisions of ACC. There was no difference in the *BP*_P_ values among subdivisions of ACC (F(3, 48)=1.06, p=0.375, One-way ANOVA). SP scores correlated negatively with the *BP*_P_ values in pgACC, aMCC and pMCC but not in subcallosal cortex (pgACC, r_s_ = -0.59, p = 0.037; aMCC, r_s_ = -0.60, p = 0.034; pMCC, r_s_ = - 0.66, p = 0.017; subcallosal cortex, r_s_ = -0.46, p = 0.117, Spearman’s rank test with FDR correction; Figure 2).

**Figure 2.**
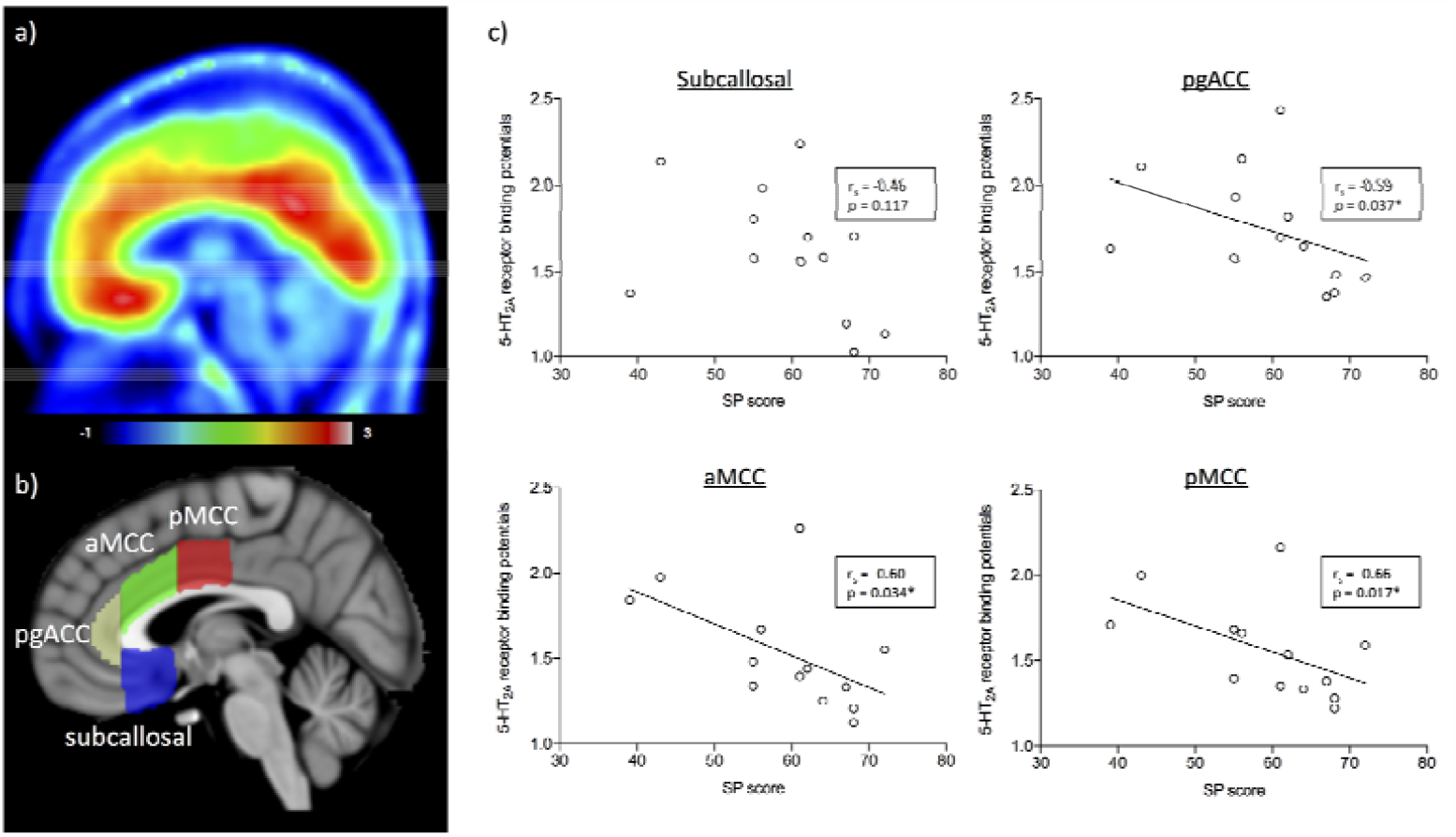
a) Mean parametric image of 5-HT_2A_ receptor binding of [^18^F]altanserin PET, shown in sagittal view. b) Subdivisions of ACC, overlaid on sagittal T1 MRI template. c) Plot graph of sensitivity to punishment (SP) score and regional 5-HT_2A_ receptor binding potentials (*BP*_P_). SP was negatively correlated with 5-HT_2A_ receptor *BP*_P_ in pgACC, aMCC and pMCC, whereas no such association was detected in subcallosal region (*false discovery rate corrected p < 0.05). Spearman’s rank test was used. pgACC, pregenual anterior cingulate cortex; aMCC, anterior midcingulate cortex; pMCC, posterior midcingulate cortex

### Association between 5-HT_2A_ receptor availability and functional connectivity

Seed-based functional connectivity analyses were performed for above three ACC subregions (Figure 3, Table 1) to explore functional connectivity that correlated with the local BP_P_ value. The *BP*_P_ values in pgACC were negatively correlated with the functional connectivity between pgACC and clusters in left lateral occipital cortex and right lingual gyrus (Figure 3a). The *BP*_P_ values in aMCC were positively correlated with the functional connectivity between aMCC and left middle frontal gyrus (MFG) (Figure 3b). The *BP*_P_ values in pMCC were positively correlated with the functional connectivity between pMCC and clusters in right inferior frontal gyrus, left precentral gyrus, left supramarginal gyrus and left angular gyrus (Figure 3c).

**Table 1.**
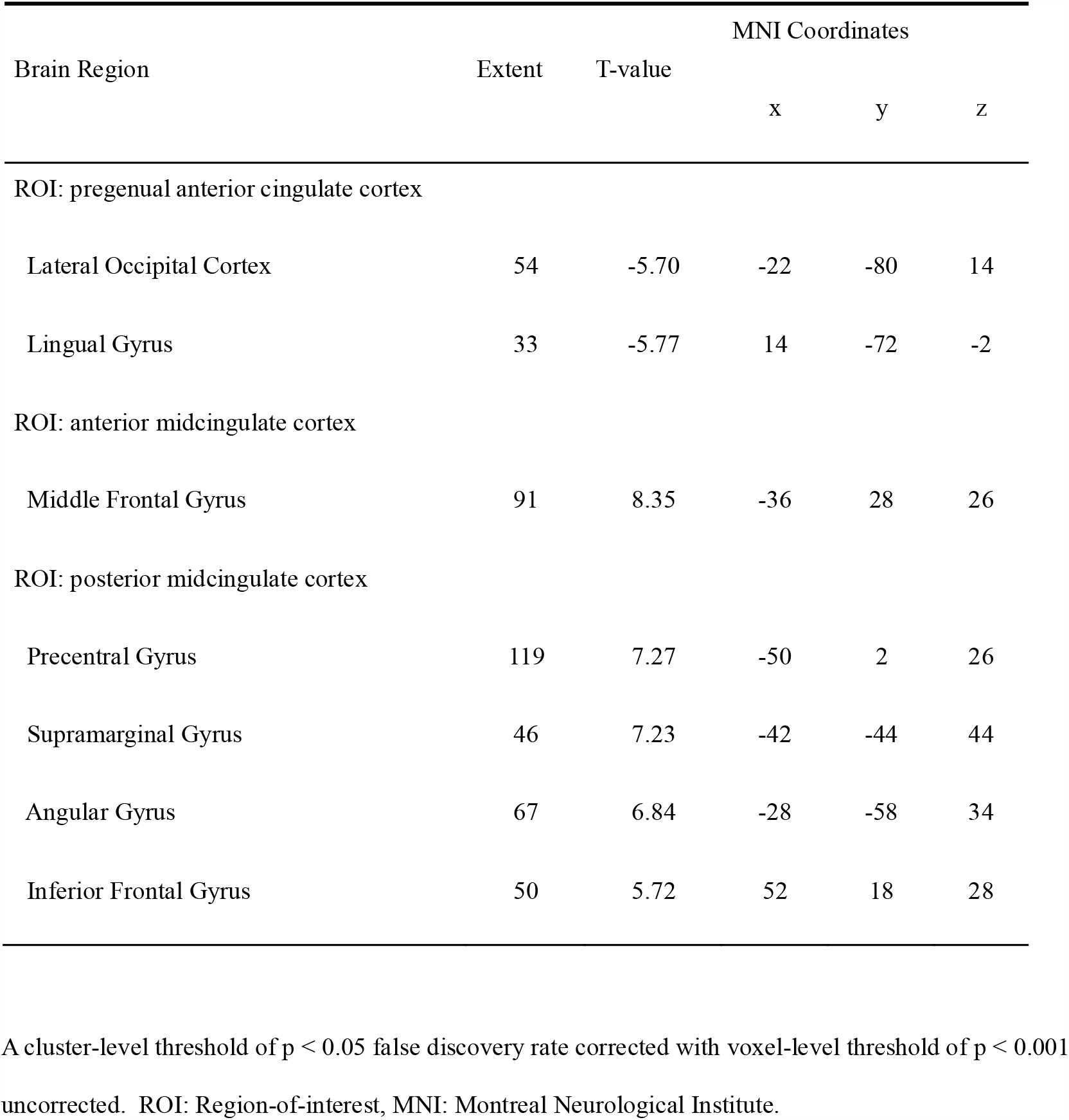
Coordinates of functional connectivity that correlated with 5-HT_2A_ receptor binding potentials in each anterior cingulate cortex subregion.

| Brain Region | Extent | T-value | MNI Coordinates |  |  |
| --- | --- | --- | --- | --- | --- |
|  |  |  | x | y | z |
| ROI: pregenual anterior cingulate cortex |  |  |  |  |  |
| Lateral Occipital Cortex | 54 | -5.70 | -22 | -80 | 14 |
| Lingual Gyrus | 33 | -5.77 | 14 | -72 | -2 |
| ROI: anterior midcingulate cortex |  |  |  |  |  |
| Middle Frontal Gyrus | 91 | 8.35 | -36 | 28 | 26 |
| ROI: posterior midcingulate cortex |  |  |  |  |  |
| Precentral Gyrus | 119 | 7.27 | -50 | 2 | 26 |
| Supramarginal Gyrus | 46 | 7.23 | -42 | -44 | 44 |
| Angular Gyrus | 67 | 6.84 | -28 | -58 | 34 |
| Inferior Frontal Gyrus | 50 | 5.72 | 52 | 18 | 28 |

A cluster-level threshold of $p < 0.05$ false discovery rate corrected with voxel-level threshold of $p < 0.001$ uncorrected. ROI: Region-of-interest, MNI: Montreal Neurological Institute.

**Figure 3.**
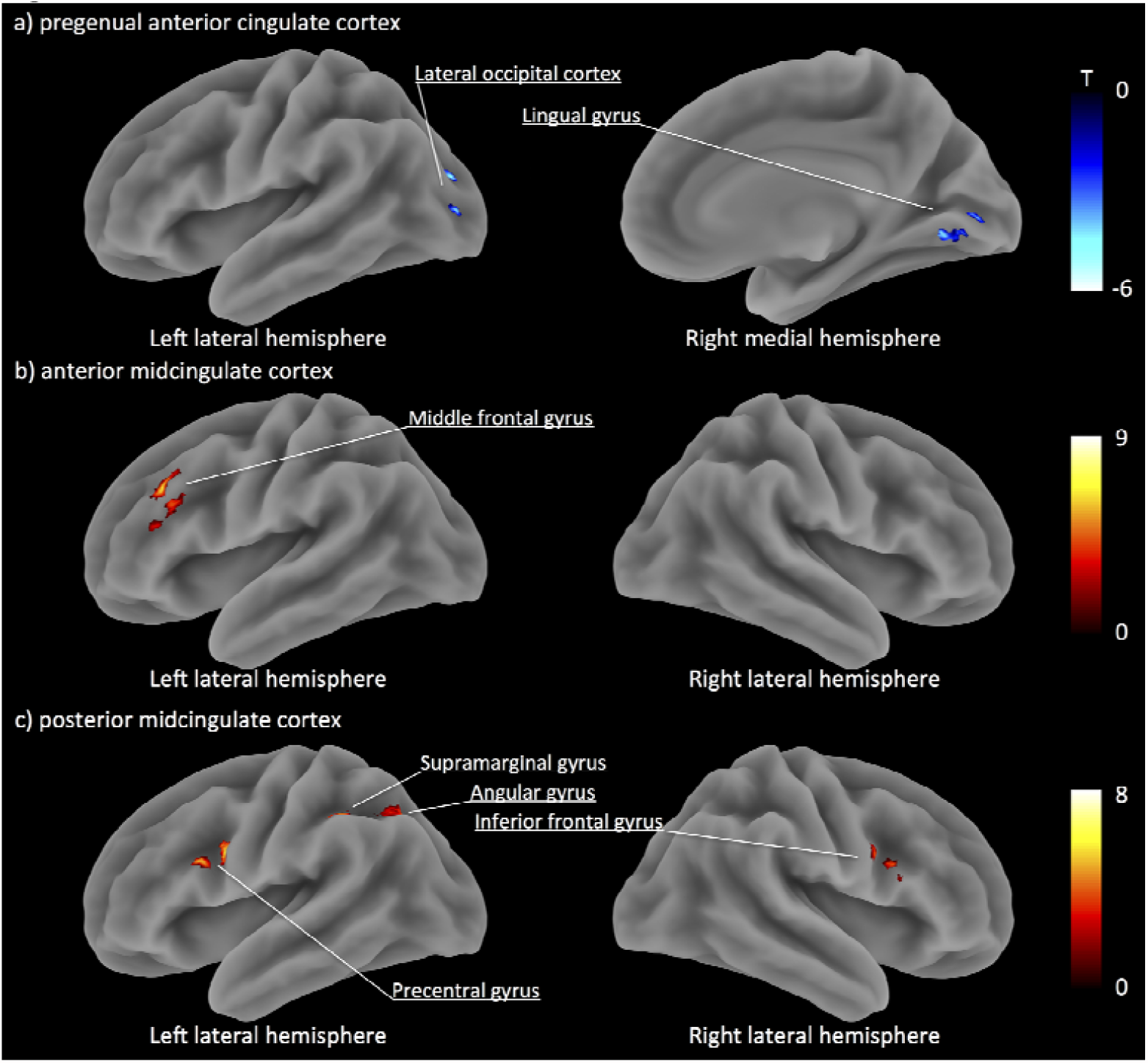
Surface rendered images of functional connectivity associated with 5-HT_2A_ receptor binding potentials in each anterior cingulate cortex subregion. a) The functional connectivity of the pregenual anterior cingulate cortex. b) The functional connectivity of the anterior midcingulate cortex. c) The functional connectivity of the posterior midcingulate cortex. Shown clusters remained after a threshold of cluster-level p < 0.05 false discovery rate corrected and voxel-level p < 0.001 uncorrected for multiple comparisons. Clusters were surface-rendered onto a brain template. Color bar represents T-value; negative correlations as blue-purple, positive correlations as red-yellow.

### Functional connectivity related to behavioral inhibition system

Correlation analyses between these specific functional connectivities and SP were carried out. SP was negatively correlated with the functional connectivity between aMCC and left MFG (r_s_ = -0.67, p = 0.014, Spearman’s rank test; Table 2).

**Table 2.** Correlations between sensitivity to punishment score and functional connectivity of each subregion of anterior cingulate cortex.

| | $r_s$ | p value |
| --- | --- | --- |
| ROI: pregenual anterior cingulate cortex (pgACC) |  |  |
| pgACC - left lateral occipital cortex | 0.52 | 0.069 |
| pgACC - right lingual gyrus | 0.44 | 0.135 |
| ROI: anterior midcingulate cortex (aMCC) |  |  |
| aMCC - left middle frontal gyrus | -0.67 | 0.014* |
| ROI: posterior mid cingulate cortex (pMCC) |  |  |
| pMCC - left precentral gyrus | -0.46 | 0.112 |
| pMCC - left angular gyrus | -0.39 | 0.189 |
| pMCC - right inferior frontal gyrus | -0.35 | 0.243 |
| pMCC - left supramarginal gyrus | -0.41 | 0.166 |

$r_s$ : Spearman's rank correlation coefficient
\* $p < 0.05$
ROI: Region-of-interest

### Mediation analysis

Finally, we examined whether functional connectivity between aMCC and left MFC serve as a potential mediator of the link between SP scores and 5-HT_2A_ receptor availability. We tested two possible models: 1) 5-HT_2A_ receptor availability affects functional connectivity, which in turn affects SP; 2) SP affects functional connectivity, which in turn affects 5-HT_2A_ receptor availability. Mediation analyses supported the latter model, indicating that the total indirect effect of SP scores on the *BP*_P_ values via functional connectivity was significant (BCa CI: -0.044 to -0.007; p < 0.05). In sum, these results imply that the functional connectivity between aMCC and left MFG serves as an important role in linking BIS with the 5-HT_2A_ receptor availability.

## Discussion

This study investigated whether the individual variations in 5-HT_2A_ receptor availability contributes to BIS and which brain networks were specifically involved. BIS correlated negatively with 5-HT_2A_ receptor availability in ACC, and the association between BIS and 5-HT_2A_ receptor availability was accounted for by the functional connectivity between aMCC and left MFG.

Our findings indicate the role of serotonergic neurotransmission in ACC, as was previously linked with BIS personality and aversive anticipation. Specifically, high BIS individuals showed reduced levels of serotonin 5-HT_2A_ receptor availability in ACC. The 5-HT_2A_ receptor is known to be related to psychiatric symptoms, such as anxiety and depression, as well as hallucinations in schizophrenia (Quednow, Geyer, & Halberstadt, 2009) and Parkinson’s disease (Ballanger et al., 2010). In recent genetic studies, polymorphism of the 5-HT_2A_ gene has been frequently reported in depression and schizophrenia (Gu et al., 2013; Tan et al., 2014; Zhao et al., 2014). For example, single nucleotide polymorphism of 5-HT_2A_ gene was associated with pathological gambling and suicide in depressed patients (Arias et al., 2001; Wilson, da Silva Lobo, Tavares, Gentil, & Vallada, 2013). Furthermore, in PET studies, while suicide victims had a high density of 5-HT_2A_ receptors in prefrontal cortex (Du, Faludi, Palkovits, Bakish, & Hrdina, 2001), treatment-resistant depressed patients displayed lower 5-HT_2A_ receptor binding in dorsal prefrontal cortex and ACC (Baeken, De Raedt, & Bossuyt, 2012). These contradictory findings represent the activation and the inhibition of impulsivity (Fineberg et al., 2010), and the current finding supports to the latter. Animal experiments have also shown that blockage of the 5-HT_2A_ receptor in medial prefrontal cortex suppressed impulsive behavior (Fink et al., 2015). Taken together, present result indicates that high BIS individuals showed stronger depressive hopelessness and facilitates lower 5-HT_2A_ receptor availability in ACC. This may reflect the specific involvement of inhibitory control associated with some aspects of depressive symptoms.

This study newly identified the functional connectivity associated with 5-HT_2A_ receptor availability in ACC subregions (pgACC, aMCC, pMCC), and the mediation test revealed that aMCC-MFG functional connectivity contributed to the link between BIS and 5-HT_2A_ receptor availability. ACC has been consistently linked with cognitive function, emotion processing, and the autonomic nervous system (Bush, Luu, & Posner, 2000; Critchley, Mathias, & Dolan, 2001), which is divided into 4 subregions (Vogt, Berger, & Derbyshire, 2003). The pgACC, the ventral part of ACC, is involved in assessing emotional and motivational information. The pMCC and aMCC are part of the dorsal ACC, mainly involved in cognitive controls. Although the implications of these functional connectivities are still to be examined in the future, the current findings of pgACC functional connectivity with visual areas and pMCC functional connectivity with fronto-parietal attention networks may suggest serotonergic modulation of motivational visual processing and attentional control, respectively.

The aMCC represents a hub where information about punishment and negative feedback, such as pain, is monitored, triggering control signals and/or selective attention generated in dorsolateral prefrontal cortex (DLPFC) (MacDonald, Cohen, Stenger, & Carter, 2000; Miller & Cohen, 2001; Shackman et al., 2011; Walsh, Buonocore, Carter, & Mangun, 2011; Yeung et al., 2004). Other studies have also shown that aMCC is anatomically connected with DLPFC in monkeys (Morecraft & Tanji, 2009), and that the functional connectivity between these two regions is correlated with working memory demand according to task-based fMRI (Osaka et al., 2004). Consistent with these previous studies, our finding of the functional connectivity between aMCC and left MFG, a part of DLPFC, possibly reflects the cognitive control associated with negative information processing, and in particular, serves a mechanistic role in linking BIS and serotonergic neurotransmission.

It is puzzling that the direction of the path was found to be from the psychological trait to the molecular system, not vice versa. The mediation test taps on the mathematical linkage rather than on a biological one. Serotonergic stimulation by physiological and acute or chronic pharmacological manner may exhibit differently in the brain function. To further clarify this question, acute and chronic serotonergic intervention may alter both BIS and the functional connectivity between aMCC and left MFG, which shall be left for the future investigations.

There are several limitations to this study. The first is that our sample size is comparatively small. Although we have set a stringent statistical threshold, future study with a larger sample size will be required to replicate the current findings. Second, females were not included in the present study. As estrogen promotes 5-HT synthesis and menstrual cycle, it influences 5-HT_2A_ receptor binding in women (Wihlbäck et al., 2004), thereby we exclusively included male subjects in the current study. Considering that emotional reactions differ between genders, it may be interesting to explore the similarities and differences between male and female subjects in the future. In addition, the enrolled subjects were all Japanese. Previous behavioral studies have indicated that Japanese are motivated more by negative feedbacks than by positive ones (Diener, Oishi, & Lucas, 2003; Heine et al., 2001); thus, our results might be biased in this regard. Lastly, functional connectivity only accounts for a linear association between two brain regions. Whole brain networks and anatomical connectivities were not examined in the present study and should be addressed in the future studies.

## Conclusions

In summary, this multimodal neuroimaging study provides novel evidence of the relationship between the behavioral inhibition and the serotonergic function, which is mediated by the functional connectivity between aMCC and left MFG, known as a cognitive control network. The link obtained in the current study may be tested by interventional studies using drugs which modulate serotonergic neurotransmission to draw out any biological causal relationships. From the basis from the current findings of healthy subjects, future studies of patients with anxiety, depression, and pain disorder symptoms may shed a light on, further understanding of 5-HT_2A_ receptor function and symptoms associated with behavioral inhibition.

## Conflict of interests

On behalf of all authors, the corresponding author states that there is no conflict of interest.

## Ethical Approval

The study was approved by the Ethics and Radiation Safety Committee of the National Institute of Radiological Sciences in accordance with the ethical standards laid down in the 1964 Declaration of Helsinki and its later amendments.

## Consent to Participate

All participants provided written informed consent before participating in the study.

## Consent to Publish

All authors provided their consent to publish this manuscreipt.

## Authors Contributions

M.Y., Y.K, T.S designed the study; M.Y., Y.K., K.T., T.I., K.Y., H.H., K.Kawamura, M.Z. and H.I. conducted the experiment; K.Kojima., M.Y., Y.K., C.S. and Y.I. analyzed data; K.Kojima, M.Y., S.H., Y.K., S.K., M.H. and T.S. wrote the paper.

## Funding

This work was supported in part by the Precursory Research for Embryonic Science and Technology, Japan Science and Technology Agency; by the Naito Foundation; by AMED under Grant Number dm0207007 and dm0107094; and by JSPS KAKENHI Grant Number 17H02173 and 20H05711.

## Competing Interests

None of the authors have conflicts of interest to disclose.

## Availability of data and materials

Not applicable

## Acknowledgements

We thank K. Suzuki, S. Kawakami, and I. Kaneko for assistance as clinical research coordinators, H. Sano for assistance as an MRI technician, and members of the Clinical Imaging Team for support with PET scans. This work was supported in part by the Precursory Research for Embryonic Science and Technology, Japan Science and Technology Agency; by the Naito Foundation; by AMED under Grant Number dm0207007 and dm0107094; and by JSPS KAKENHI Grant Number 17H02173.

## Notes

**Conflict of interests** The authors declare no competing interests.

### Competing Interest Statement

The authors have declared no competing interest.

## References

Arias, B., Gasto, C., Catalan, R., Gutierrez, B., Pintor, L., & Fananas, L. (2001). The 5-HT(2A) receptor gene 102T/C polymorphism is associated with suicidal behavior in depressed patients. Am J Med Genet, 105(8), 801–804. doi:10.1002/ajmg.10099

Baeken, C., Bossuyt, A., & De Raedt, R. (2014). Dorsal prefrontal cortical serotonin 2A receptor binding indices are differentially related to individual scores on harm avoidance. Psychiatry Res, 221(2), 162–168. doi:10.1016/j.pscychresns.2013.12.005

Baeken, C., De Raedt, R., & Bossuyt, A. (2012). Is treatment-resistance in unipolar melancholic depression characterized by decreased serotonin(2)A receptors in the dorsal prefrontal - anterior cingulate cortex? Neuropharmacology, 62(1), 340–346. doi:10.1016/j.neuropharm.2011.07.043

Baldwin, D., & Rudge, S. (1995). The role of serotonin in depression and anxiety. Int Clin Psychopharmacol, 9 Suppl 4, 41–45. doi:10.1097/00004850-199501004-00006

Ballanger, B., Strafella, A. P., van Eimeren, T., Zurowski, M., Rusjan, P. M., Houle, S., & Fox, S. H. (2010). Serotonin 2A receptors and visual hallucinations in Parkinson disease. Arch Neurol, 67(4), 416–421. doi:10.1001/archneurol.2010.35

Barros-Loscertales, A., Meseguer, V., Sanjuan, A., Belloch, V., Parcet, M. A., Torrubia, R., & Avila, C. (2006). Behavioral Inhibition System activity is associated with increased amygdala and hippocampal gray matter volume: A voxel-based morphometry study. Neuroimage, 33(3), 1011–1015. doi:10.1016/j.neuroimage.2006.07.025

Beaver, J. D., Lawrence, A. D., Passamonti, L., & Calder, A. J. (2008). Appetitive motivation predicts the neural response to facial signals of aggression. J Neurosci, 28(11), 2719–2725. doi:10.1523/JNEUROSCI.0033-08.2008

Beck, A. T., Weissman, A., Lester, D., & Trexler, L. (1974). The measurement of pessimism: the hopelessness scale. J Consult Clin Psychol, 42(6), 861–865. doi:10.1037/h0037562

Bijttebier, P., Beck, I., Claes, L., & Vandereycken, W. (2009). Gray’s Reinforcement Sensitivity Theory as a framework for research on personality-psychopathology associations. Clin Psychol Rev, 29(5), 421–430. doi:10.1016/j.cpr.2009.04.002

Bush, G., Luu, P., & Posner, M. I. (2000). Cognitive and emotional influences in anterior cingulate cortex. Trends Cogn Sci, 4(6), 215–222. doi:10.1016/s1364-6613(00)01483-2

Cherbuin, N., Windsor, T. D., Anstey, K. J., Maller, J. J., Meslin, C., & Sachdev, P. S. (2008). Hippocampal volume is positively associated with behavioural inhibition (BIS) in a large community-based sample of mid-life adults: the PATH through life study. Soc Cogn Affect Neurosci, 3(3), 262–269. doi:10.1093/scan/nsn018

Cloninger, C. R. (1987). A systematic method for clinical description and classification of personality variants. A proposal. Arch Gen Psychiatry, 44(6), 573–588. doi:10.1001/archpsyc.1987.01800180093014

Corr, P. J. (2002). J. A. Gray’s reinforcement sensitivity theory: tests of the joint subsystems hypothesis of anxiety and impulsivity. Personality and Individual Differences, 33(4), 511–532. doi:10.1016/S0191-8869(01)00170-2

Corr, P. J. (2004). Reinforcement sensitivity theory and personality. Neurosci Biobehav Rev, 28(3), 317–332. doi:10.1016/j.neubiorev.2004.01.005

Critchley, H. D., Mathias, C. J., & Dolan, R. J. (2001). Neuroanatomical basis for first- and second-order representations of bodily states. Nat Neurosci, 4(2), 207–212. doi:10.1038/84048

Davidson, R. J. (1994). Cerebral Asymmetry, Emotion, and Affective Style. In R.J. Davidson & K. Hugdahl (Eds.), Brain Asymmetry (pp. 361–387). Massachusetts: MIT Press.

De Martino, B., Camerer, C. F., & Adolphs, R. (2010). Amygdala damage eliminates monetary loss aversion. Proc Natl Acad Sci U S A, 107(8), 3788–3792. doi:10.1073/pnas.0910230107

Diener, E., Oishi, S., & Lucas, R. E. (2003). Personality, culture, and subjective well-being: emotional and cognitive evaluations of life. Annu Rev Psychol, 54, 403–425. doi:10.1146/annurev.psych.54.101601.145056

Du, L., Faludi, G., Palkovits, M., Bakish, D., & Hrdina, P. D. (2001). Serotonergic genes and suicidality. Crisis, 22(2), 54–60. doi:10.1027//0227-5910.22.2.54

Eisenberger, N. I. (2012). The pain of social disconnection: examining the shared neural underpinnings of physical and social pain. Nat Rev Neurosci, 13(6), 421–434. doi:10.1038/nrn3231

Fineberg, N. A., Potenza, M. N., Chamberlain, S. R., Berlin, H. A., Menzies, L., Bechara, A., … Hollander, E. (2010). Probing compulsive and impulsive behaviors, from animal models to endophenotypes: a narrative review. Neuropsychopharmacology, 35(3), 591–604. doi:10.1038/npp.2009.185

Fink, L. H., Anastasio, N. C., Fox, R. G., Rice, K. C., Moeller, F. G., & Cunningham, K. A. (2015). Individual Differences in Impulsive Action Reflect Variation in the Cortical Serotonin 5-HT2A Receptor System. Neuropsychopharmacology, 40(8), 1957–1968. doi:10.1038/npp.2015.46

Fuentes, P., Barros-Loscertales, A., Bustamante, J. C., Rosell, P., Costumero, V., & Avila, C. (2012). Individual differences in the Behavioral Inhibition System are associated with orbitofrontal cortex and precuneus gray matter volume. Cogn Affect Behav Neurosci, 12(3), 491–498. doi:10.3758/s13415-012-0099-5

Gray, J. A. (1982). The neuropsychology of anxiety: An enquiry into the functions of the septo-hippocampal system. . New York: Oxford University Press,.

Gray, J. A., & McNaughton, N. (2000). The neuropsychology of anxiety: An enquiry into the functions of the septo-hippocampal system. (2 ed. Vol. 33). New York: Oxford University Press,.

Gu, L., Long, J., Yan, Y., Chen, Q., Pan, R., Xie, X., … Su, L. (2013). HTR2A-1438A/G polymorphism influences the risk of schizophrenia but not bipolar disorder or major depressive disorder: a meta-analysis. J Neurosci Res, 91(5), 623–633. doi:10.1002/jnr.23180

Hahn, T., Dresler, T., Pyka, M., Notebaert, K., & Fallgatter, A. J. (2013). Local synchronization of resting-state dynamics encodes Gray’s trait Anxiety. PLoS One, 8(3), e58336. doi:10.1371/journal.pone.0058336

Hayes, D. J., & Northoff, G. (2011). Identifying a network of brain regions involved in aversion-related processing: a cross-species translational investigation. Front Integr Neurosci, 5, 49. doi:10.3389/fnint.2011.00049

Heine, S. J., Lehman, D. R., Ide, E., Leung, C., Kitayama, S., Takata, T., & Matsumoto, H. (2001). Divergent consequences of success and failure in japan and north america: an investigation of self-improving motivations and malleable selves. J Pers Soc Psychol, 81(4), 599–615. doi:10.1037/0022-3514.81.4.599

Higgins, E. T., Roney, C. J., Crowe, E., & Hymes, C. (1994). Ideal versus ought predilections for approach and avoidance: distinct self-regulatory systems. J Pers Soc Psychol, 66(2), 276–286. doi:10.1037/0022-3514.66.2.276

Innis, R. B., Cunningham, V. J., Delforge, J., Fujita, M., Gjedde, A., Gunn, R. N., … Carson, R. E. (2007). Consensus nomenclature for in vivo imaging of reversibly binding radioligands. J Cereb Blood Flow Metab, 27(9), 1533–1539. doi:10.1038/sj.jcbfm.9600493

Ishii, T., Kimura, Y., Ichise, M., Takahata, K., Kitamura, S., Moriguchi, S., … Suhara, T. (2017). Anatomical relationships between serotonin 5-HT2A and dopamine D2 receptors in living human brain. PLoS One, 12(12), e0189318. doi:10.1371/journal.pone.0189318

Jensen, M. P., Ehde, D. M., & Day, M. A. (2016). The Behavioral Activation and Inhibition Systems: Implications for Understanding and Treating Chronic Pain. J Pain, 17(5), 529 e521–529 e518. doi:10.1016/j.jpain.2016.02.001

Kim, S. H., Yoon, H., Kim, H., & Hamann, S. (2015). Individual differences in sensitivity to reward and punishment and neural activity during reward and avoidance learning. Soc Cogn Affect Neurosci, 10(9), 1219–1227. doi:10.1093/scan/nsv007

Kringelbach, M. L., & Rolls, E. T. (2004). The functional neuroanatomy of the human orbitofrontal cortex: evidence from neuroimaging and neuropsychology. Progress in Neurobiology, 72(5), 341–372. doi:10.1016/j.pneurobio.2004.03.006.

Logan, J., Fowler, J. S., Volkow, N. D., Wolf, A. P., Dewey, S. L., Schlyer, D. J., … et al. (1990). Graphical analysis of reversible radioligand binding from time-activity measurements applied to [N-11C-methyl]-(-)-cocaine PET studies in human subjects. J Cereb Blood Flow Metab, 10(5), 740–747. doi:10.1038/jcbfm.1990.127

MacDonald, A. W., 3rd, Cohen, J.D., Stenger, V. A., & Carter, C. S. (2000). Dissociating the role of the dorsolateral prefrontal and anterior cingulate cortex in cognitive control. Science, 288(5472), 1835–1838. doi:10.1126/science.288.5472.1835

Miller, E. K., & Cohen, J. D. (2001). An integrative theory of prefrontal cortex function. Annu Rev Neurosci, 24, 167–202. doi:10.1146/annurev.neuro.24.1.167

Morecraft, R. J., & Tanji, J. (2009). Cingulofrontal Interactions and the Cingulate Motor Areas. In V. A. Vogt (Ed.), Cingulate Neurobiology and Disease (pp. 114–144). New York: Oxford University Press.

Moresco, F. M., Dieci, M., Vita, A., Messa, C., Gobbo, C., Galli, L., … Fazio, F. (2002). In vivo serotonin 5HT(2A) receptor binding and personality traits in healthy subjects: a positron emission tomography study. Neuroimage, 17(3), 1470–1478. doi:10.1006/nimg.2002.1239

Nitschke, J. B., Sarinopoulos, I., Mackiewicz, K. L., Schaefer, H. S., & Davidson, R. J. (2006). Functional neuroanatomy of aversion and its anticipation. Neuroimage, 29(1), 106–116. doi:10.1016/j.neuroimage.2005.06.068

Osaka, N., Osaka, M., Kondo, H., Morishita, M., Fukuyama, H., & Shibasaki, H. (2004). The neural basis of executive function in working memory: an fMRI study based on individual differences. Neuroimage, 21(2), 623–631. doi:10.1016/j.neuroimage.2003.09.069

Price, J. C., Lopresti, B. J., Meltzer, C. C., Smith, G. S., Mason, N. S., Huang, Y., … Mathis, C. A. (2001). Analyses of [(18)F]altanserin bolus injection PET data. II: consideration of radiolabeled metabolites in humans. Synapse, 41(1), 11–21. doi:10.1002/syn.1055

Quednow, B. B., Geyer, M. A., & Halberstadt, A. L. (2009). Serotonin and Schizophrenia. In P.M. hristian & L. J. Barry (Eds.), Handbook of the Behavioral Neurobiology of Serotonin. (2 ed., Vol. 21, pp. 585–620). London: Academic Press.

Savli, M., Bauer, A., Mitterhauser, M., Ding, Y. S., Hahn, A., Kroll, T., … Lanzenberger, R. (2012). Normative database of the serotonergic system in healthy subjects using multi-tracer PET. Neuroimage, 63(1), 447–459. doi:10.1016/j.neuroimage.2012.07.001

Shackman, A. J., Salomons, T. V., Slagter, H. A., Fox, A. S., Winter, J. J., & Davidson, R. J. (2011). The integration of negative affect, pain and cognitive control in the cingulate cortex. Nat Rev Neurosci, 12(3), 154–167. doi:10.1038/nrn2994

Simon, J. J., Walther, S., Fiebach, C. J., Friederich, H. C., Stippich, C., Weisbrod, M., & Kaiser, S. (2010). Neural reward processing is modulated by approach- and avoidance-related personality traits. Neuroimage, 49(2), 1868–1874. doi:10.1016/j.neuroimage.2009.09.016

Soloff, P. H., Price, J. C., Mason, N. S., Becker, C., & Meltzer, C. C. (2010). Gender, personality, and serotonin-2A receptor binding in healthy subjects. Psychiatry Res, 181(1), 77–84. doi:10.1016/j.pscychresns.2009.08.007

Sommer, C. (2009). Serotonin in Pain and Pain Control. In P.M. Christian & L. J. Barry (Eds.), Handbook of the Behavioral Neurobiology of Serotonin. (2 ed., Vol. 21, pp. 457–471). London: Academic Press.

Spielberger, C. D., Gorsuch, R. L., Lushene, R. E., Vagg, P. R., & Jacob, G. A. (1983). Manual for the State-trait Anxiety Inventory. California: Consulting Psychologist Press.

Takahashi, Y., & Shigemasu, K. (2008). Comparison of Three Scales Measuring Individual Differences in Sensitivity to Punishment and Reward. The Japanese Journal of Personality, 17(1), 72–81. doi:10.2132/personality.17.72

Tan, J., Chen, S., Su, L., Long, J., Xie, J., Shen, T., … Gu, L. (2014). Association of the T102C polymorphism in the HTR2A gene with major depressive disorder, bipolar disorder, and schizophrenia. Am J Med Genet B Neuropsychiatr Genet, 165B(5),p438-455. doi:10.1002/ajmg.b.32248

Torrubia, R., Ávilab, C., Moltó, J., & Caseras, X. (2001). The Sensitivity to Punishment and Sensitivity to Reward Questionnaire (SPSRQ) as a measure of Gray’s anxiety and impulsivity dimensions. Personality and Individual Differences, 31(6), 837–862. doi:10.1016/S0191-8869(00)00183-5

Vogt, B. A., Berger, G. R., & Derbyshire, S. W. (2003). Structural and functional dichotomy of human midcingulate cortex. Eur J Neurosci, 18(11), 3134–3144. doi:10.1111/j.1460-9568.2003.03034.x

Walsh, B. J., Buonocore, M. H., Carter, C. S., & Mangun, G. R. (2011). Integrating conflict detection and attentional control mechanisms. J Cogn Neurosci, 23(9), 2211–2221. doi:10.1162/jocn.2010.21595

Wardak, M., Wong, K. P., Shao, W., Dahlbom, M., Kepe, V., Satyamurthy, N., … Huang, S. C. (2010). Movement correction method for human brain PET images: application to quantitative analysis of dynamic 18F-FDDNP scans. J Nucl Med, 51(2), 210–218. doi:10.2967/jnumed.109.063701

Wihlbäck, A. C., Sundstrom Poromaa, I., Bixo, M., Allard, P., Mjorndal, T., & Spigset, O. (2004). Influence of menstrual cycle on platelet serotonin uptake site and serotonin2A receptor binding. Psychoneuroendocrinology, 29(6), 757–766. doi:10.1016/S0306-4530(03)00120-3

Wilson, D., da Silva Lobo, D.S., Tavares, H., Gentil, V., & Vallada, H. (2013). Family-based association analysis of serotonin genes in pathological gambling disorder: evidence of vulnerability risk in the 5HT-2A receptor gene. J Mol Neurosci, 49(3), 550–553. doi:10.1007/s12031-012-9846-x

Wrase, J., Kahnt, T., Schlagenhauf, F., Beck, A., Cohen, M. X., Knutson, B., & Heinz, A. (2007). Different neural systems adjust motor behavior in response to reward and punishment. Neuroimage, 36(4), 1253–1262. doi:10.1016/j.neuroimage.2007.04.001

Yeung, N., Botvinick, M. M., & Cohen, J. D. (2004). The neural basis of error detection: conflict monitoring and the error-related negativity. Psychol Rev, 111(4), 931–959. doi:10.1037/0033-295X.111.4.931

Zhao, X., Sun, L., Sun, Y. H., Ren, C., Chen, J., Wu, Z. Q., … Lv, X. L. (2014). Association of HTR2A T102C and A-1438G polymorphisms with susceptibility to major depressive disorder: a meta-analysis. Neurol Sci, 35(12), 1857–1866. doi:10.1007/s10072-014-1970-7

